# Vendor-neutral sequences and fully transparent workflows improve inter-vendor reproducibility of quantitative MRI

**DOI:** 10.1101/2021.12.27.474259

**Authors:** Agah Karakuzu, Labonny Biswas, Julien Cohen-Adad, Nikola Stikov

## Abstract

**Purpose:** We developed an end-to-end workflow that starts with a vendor-neutral acquisition and tested the hypothesis that vendor-neutral sequences decrease inter-vendor variability of T1, MTR and MTsat measurements.

**Methods:** We developed and deployed a vendor-neutral 3D spoiled gradient-echo (SPGR) sequence on three clinical scanners by two MRI vendors. We then acquired T1 maps on the ISMRM-NIST system phantom, as well as T1, MTR and MTsat maps in three healthy participants. We performed hierarchical shift function analysis in vivo to characterize the differences between scanners when the vendor-neutral sequence is used instead of commercial vendor implementations. Inter-vendor deviations were compared for statistical significance to test the hypothesis.

**Results:** In the phantom, the vendor-neutral sequence reduced inter-vendor differences from 8 - 19.4% to 0.2 - 5% with an overall accuracy improvement, reducing ground truth T1 deviations from 7 - 11% to 0.2 - 4%. In vivo we found that the variability between vendors is significantly reduced (p = 0.015) for all maps (T1, MTR and MTsat) using the vendor-neutral sequence.

**Conclusion:** We conclude that vendor-neutral workflows are feasible and compatible with clinical MRI scanners. The significant reduction of inter-vendor variability using vendor-neutral sequences has important implications for qMRI research and for the reliability of multicenter clinical trials.

## Introduction

As the invention of MRI approaches its 50th anniversary^1^, the notion of image acquisition has almost become synonymous with data collection. A major driving force in the transformation of MR images from mere pictures into mineable data^2^ is attributing physiologically relevant physical parameters to the voxels, namely quantitative MRI (qMRI). MRI is not a quantitative measurement device by design. Nonetheless, systematic manipulation of effective micrometer-level MRI parameters via specialized acquisition methods, followed by fitting the resulting data to a signal representation or a biophysical model^3^, can yield parametric maps, turning scanners into quantitative diagnostic tools. Despite being as old as MRI itself, most of the qMRI methods have not succeeded to find widespread use in the clinic, at least in part due to a major multicenter reproducibility challenge.

The introduction is organized around two problems hampering multicenter reproducibility of qMRI, which this study seeks to address:

1. Lack of transparency and multicenter consistency in vendor implementations of pulse sequences that are commonly used in qMRI
2. Technical roadblocks in the way of deploying a standardized pulse sequence along with a unified user interface to multiple imaging sites

T1 relaxometry is a clear example of how availability, transparency and multicenter consistency of pulse sequences influence multicenter reproducibility. Several methods such as inversion-recovery spin-echo (IR-SE)^4^, variable flip angle (VFA-T1)^5^, Look-Locker IR^6^ and magnetization-prepared two rapid-echoes (MP2RAGE)^7^ have gained popularity in MRI research. Although measured T1 values can exhibit up to 30% inter-sequence variability in the same scan session for the same participant^8^, a selected T1 relaxometry method is much more reliable within-site^9,10^. As for the multicenter stability, MP2RAGE appears to be a promising T1 mapping method at 7T with a single vendor considered^11^. On the other hand, substantial multicenter variability is reported for another popular whole-brain imaging method VFA-T1, both in-vivo and in phantoms^12,13^. Several factors contribute to the variability of the VFA-T1 measurement, including B1 field inhomogeneity^8^, incomplete spoiling^14^, sequence parameters and bore temperature^13^, and uncontrolled magnetization transfer (MT) effects^15^. Because of all these diverse confounders of T1 stability, the healthy range of in-vivo T1 values at 3T remains elusive^16,17^. This constitutes a critical problem for the potential use of T1 relaxometry in clinics.

Considerable amount of research has focused on the measurement bias due to acquisition-related imperfections. However, the reproducibility of the developed techniques is often hindered by *problem 1.* For example, a simple yet powerful B1 correction framework for VFA-T1 has been established^18^, but such methods are typically not available in commercial systems, or the available ones vary across vendors. This not only imposes a practical challenge in evaluating the reliability of VFA-T1 measurements across vendors^13,19^, but the differences between vendor-native B1 mapping methods can aggravate the instability^20^. Another example is the spoiling gradient area and RF spoiling phase increment in the commercial implementations of spoiled gradient-echo (SPGR) sequences. Both parameters determine the accuracy of VFA-T1 mapping. However, vendors are known to set different defaults for these parameters, rendering some of them unfit for this application^14^. Similarly, fundamental properties of the excitation pulse (e.g., pulse shape, time-bandwidth product, duration) are not disclosed and it is not known how these properties are adjusted under different SAR requirements. To achieve a standardized SPGR acquisition for T1 mapping, such parameter configurations should be disclosed, made accessible and standardized across scanners for eliminating systematic biases. Recently, Gracien et al. showed a successful example of how this solution can reduce systematic biases in relaxometry mapping between two different scanner models from the same vendor^21^.

Addressing inadequacies of model assumptions constitutes another solution toward improving the reliability of qMRI methods^3^. For example, balancing the total amount of RF power deposited by each run of a VFA acquisition, Teixeira et al. enforced two-pool MT systems to behave like a single pool MT system^15^. They showed that the measurement reliability increases by making the single pool assumption valid through controlled saturation of MT. Although this technique holds important implications for multicenter reproducibility of qMRI, deploying it to multiple sites is not a straightforward process. Moreover, proprietary programming libraries of different manufacturers may not allow identical implementations, exemplifying the constraints imposed by *problem 2.* Another model-related improvement has been recently introduced to reduce the B1 dependency of MTsat maps by replacing the fixed empirical B1 correction factor of the MT saturation-index (MTsat)^22^ with a correction factor map^23^. The proposed methodology requires the details of the saturation and excitation pulses (e.g., shape, offset, duration, etc.) as the correction framework is simulation-based. From the standpoint of *problem 1,* such information is not easily accessible in the stock sequence, so the correction cannot be applied. From the perspective of *problem 2,* deploying sequences in multiple centers with known saturation and excitation pulse parameters may not be realistic due to vendor restrictions. Even though both studies made their code publicly available to facilitate the reproducibility of their work^24^, black-box vendor strategies thwart these valuable efforts.

Fortunately, there are several open-source pulse sequence development platforms to contend with *problem 2*^25–31^. These platforms can interpret and translate the same sequence logic for multiple vendors, considerably reducing multi-center development efforts and minimizing implementation variability. Another advantage of these tools is to attract community-driven development. For example, Pulseq has received considerable community attention to motivate the development of sequences in Python^28^, or even going beyond code to graphically assemble^32^ Pulseq descriptions using Pulseq-GPI^33^. Currently, Pulseq can be operated on two major clinical scanners (Siemens and GE) and three pre-clinical scanner platforms. There is recent literature showing the feasibility of Pulseq for performing multicenter qMRI studies. For example, a standardized chemical exchange saturation (CEST) protocol has been developed and deployed on three Siemens scanners, where two of the systems had different vendor software versions^34^. Results by Herz et al. showed multicenter consistency for an advanced CEST method, which has been made publicly available for both Python and Matlab users. Another recent Pulseq study performed inversion-recovery T1 mapping and multi-echo spin-echo T2 mapping on two Siemens scanners at 1.5 and 3T in phantom^35^. In that study the reference T1 mapping method^36^ accurately estimated T1 values within an 8% error band, whereas the T2 accuracy was slightly reduced. Taken together, these studies reveal the vital role of vendor-neutral pulse sequences in standardizing qMRI across centers. However, whether a vendor-neutral approach can improve quantitative agreement between scanners from different vendors has remained an open question.

The focus of earlier open-source pulse sequence platforms was providing a rapid and unified prototyping framework for facilitating interoperability, so some of the most adjusted scan parameters (e.g., field of view) had to be fixed once the sequence was downloaded to the scanner. More recent solutions such as GammaStar^31^ can remove this limitation by enabling user interaction trough the vendor’s user interface to modify fundamental protocol settings during the imaging session. Offering a more complete solution to *problem 2* through on-the-fly sequence updates, GammaStar eases the collaborative sequence development process by providing a web-based interface. Although such technical improvements reduce the barrier to entry for free sequence development, exchange and standardization, the validation aspect of open-source sequences has remained elusive. Recently, Tong et al. (2021) proposed a framework for testing, documenting and sharing open-source pulse sequences^35^, which adds an important missing piece to the community-driven MRI development puzzle.

RTHawk^37^ is another vendor-neutral solution, which is a proprietary platform for MRI software development. As it is utilizing the same infrastructure as an FDA approved (510(k), No: K1833274) cardiac imaging platform, it ensures operation within MRI hardware and safety limits. Unlike the above-mentioned solutions, RTHawk provides a remote procedure call (RPC) server that replaces the vendor’s pulse sequence controller to orchestrate vendor-specific low-level hardware instructions. The RPC pulse sequence server receives control commands and relevant sequence components (i.e., RF and gradient waveforms, ADC and timing events designed in SpinBench, as shown in Fig. 1c) directly from a standalone Ubuntu workstation connected to the scanner network (Fig. 1a). This gives the flexibility to issue synchronous or asynchronous updates to a sequence in real-time, such as scaling/replacing waveforms between TRs or changing the volume prescription. As the sequence control manager is decoupled from the vendor’s workstation, RTHawk makes it possible to develop a vendor-neutral unified user interface (UUI) per application (Fig. 1b). In addition, the collected raw data is streamed over to the standalone Ubuntu workstation through a real-time transport protocol (RTP). The RTP data manager enables adding or changing the metadata associated with each observation, which enables exporting raw and reconstructed images in community data standards (Fig. 1d).

**Figure 1.**
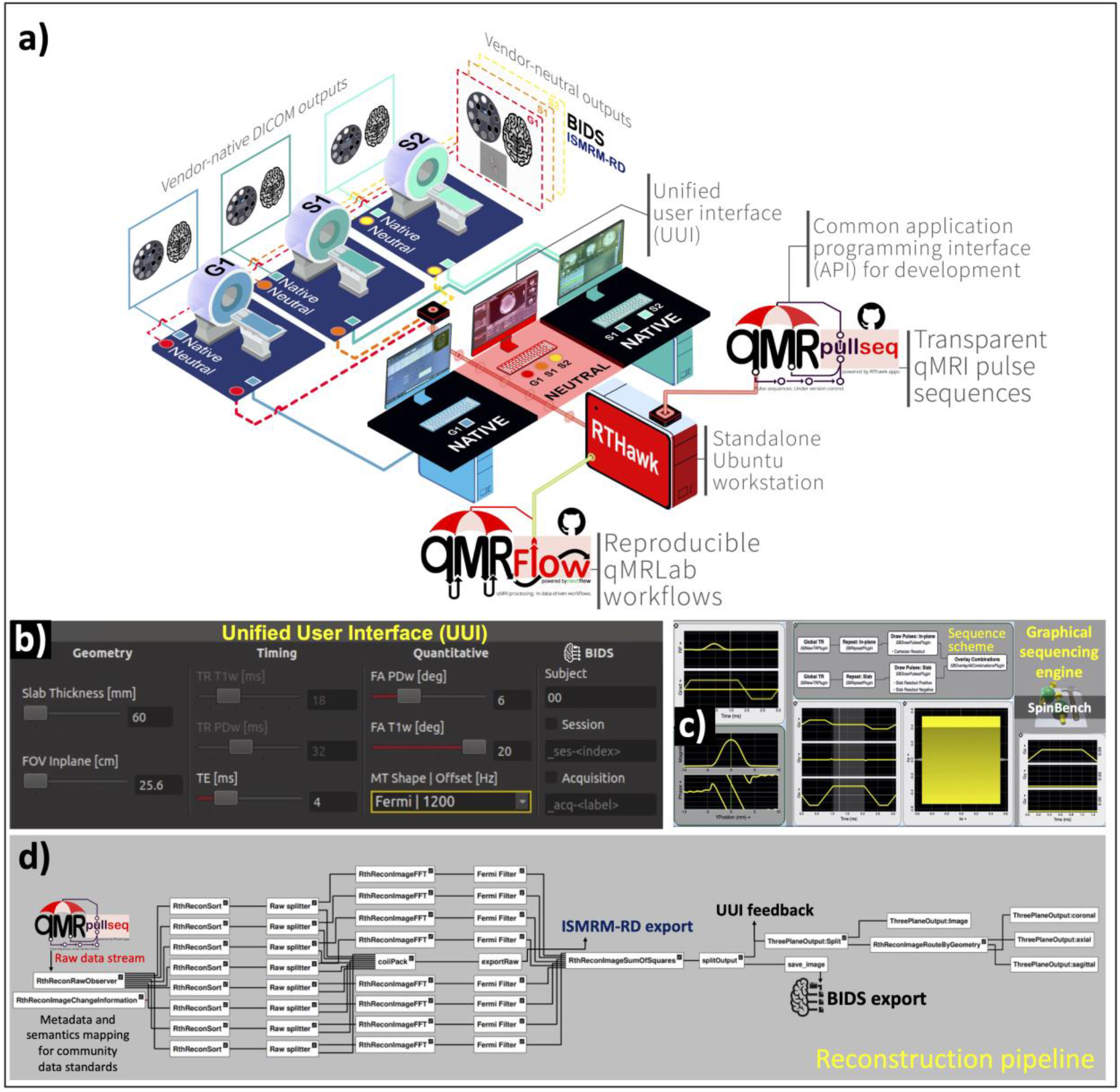
Schematic illustration of the experimental design for multicenter data collection using vendor-native and vendor-neutral pulse sequences and pulse sequence development components: **a)** 3 MRI systems are located at 2 different sites and are labeled G1 (GE 750w), S1 (Siemens Prisma) and S2 (Siemens Skyra). Vendor “Native” systems export data in the DICOM format. The proposed vendor-agnostic “Neutral” system can export a complete set of reconstructed images in BIDS and the k-space data in ISMRM-RD format, synchronized across MRI systems. Connecting to the MRI system(s) over the local network, RTHawk (red workstation) can play open-source qMRI pulse sequences under version control (qMRPullseq). All the sequences are publicly available at https://github.com/qmrlab/pulse_sequences. Fully containerized qMRFlow data-driven pipelines can connect to the scanner data stream for post-processing on the RTHawk workstation (red workstation). The same pipelines can be reproduced on a local computer, supercomputing clusters or in the cloud. **b)** The acquisitions are controlled using a unified user interface (UUI), providing a consistent user experience across vendors. **c)** RF and gradient waveform stub blocks together with the readout logic are developed using SpinBench. **d)** RTHawk reconstruction pipeline nodes are illustrated for an 8-channel receiver, also indicating how raw and reconstructed data are exported and forwarded to the display tools for on-site visualization.

Aside from vendor-neutral experiments, researchers looked at improving qMRI stability by customizing vendor-native implementations and equalizing parameters to the utmost extent possible^21,38,39^. Nevertheless, downstream data harmonization methods were still needed to correct for certain inter-vendor differences^39^, or some of the bias could not be removed altogether^38^. This is because vendor-native sequence customization may not offer a standard qMRI protocol, even for scanners with comparable hardware specs, as the selection of sequence design elements is exclusive to each manufacturer.

In this study, we test the hypothesis that vendor-neutral sequences reduce inter-vendor variability of T1, MTR and MTsat measurements. To test this hypothesis, we developed an end-to-end solution starting with a pulse sequence developed on RTHawk, followed by a fully transparent qMRLab workflow. We compared vendor-native T1, MTR and MTsat maps^40^ with those obtained using the developed vendor-neutral sequence (VENUS) workflow in three healthy participants, across three different scanners models from two manufacturers at 3T.

## Methods

### Vendor-neutral pulse sequence development

We deployed vendor-neutral pulse sequences developed in RTHawk v3.0.0 (rc4-28-ge3540dda19) (HeartVista Inc., CA, USA) on three 3T systems: (G1) GE Discovery 750 software version DV25 (R02_1549.b) (GE Healthcare, Milwaukee, MI, USA), (S1) Siemens Prisma software version VE11C (N4_LATEST_20160120) (Siemens Healthineers, Erlangen, Germany) and (S2) Siemens Skyra with the same software version as (ii). Throughout the rest of this article, these scanners will be referred to as G1, S1 and S2, respectively. Fig.-1a illustrates the hardware and software components of the experimental setup.

### General design considerations

All vendor-neutral protocols were based on a 3D SPGR pulse sequence^41^, with the RF, gradient waveforms, and the readout scheme developed as independent sequence blocks in SpinBench-v2.5.2 (Fig. 1c). To modify these sequence blocks, an RTHawk application and an additional UUI were developed for quantitative imaging, allowing the user to manage relevant acquisition parameters (e.g., FA, TR, and MT pulse for MTsat) from one simple panel that is vendor-neutral (Fig. 1b). Identical scan geometry and pre-acquisition settings were transferred between each individual acquisition. To avoid signal clipping, the highest SNR acquisition (i.e., T1w acquisition of the MTsat protocol) were run first. A simple sum-of-squares multi-coil reconstruction was developed with a Fermi filter (transition width = 0.01, radius = 0.48, both expressed as a proportion of the FOV) (Fig. 1d).

All the metadata annotations, accumulation logic of the collected data and naming of the exported images were designed according to the community data standards: ISMRM-RD^42^ for the k-space data and the Brain Imaging Data Structure (BIDS) for the reconstructed images^43,44^.

### The vendor-neutral protocol

A slab-selective (thickness = 50mm, gradient net area = 4.24 cyc/thickness) SINC excitation pulse (time-bandwidth product (TΔf) = 8, duration = 1.03ms, Hanning windowed) was implemented with a quadratic phase increment of 117° for RF spoiling. This was followed by a fully-sampled 3D cartesian readout. The default geometry properties were 256×256 acquisition matrix, 25.6 cm FOV and 20 partitions in the slab-selection direction, yielding 1×1×3 mm resolution. The readout gradient had a rewinder lobe with 2 cyc/pixel net area and was followed by a spoiling gradient with an area of 40 mT·ms/m.

For the magnetization transfer (MT) saturation, a Fermi pulse (duration = 12ms, B1rms = 3.64 μT, frequency offset = 1.2kHz, transition width = 0.35, max B1 = 5μT, pulse angle= 490°) was designed as an optional block. A loop command was defined for the sequence to iterate through three sets of parameters (i.e., MT, FA and TR), defined by the user in the UUI for a complete MTsat protocol.

From this protocol we acquired three images: (i) PD-weighted SPGR with no MT, FA = 6° and TR = 32ms (ii) MT-weighted SPGR with MT, FA = 6° and TR = 32ms (iii) T1-weighted SPGR without MT, FA = 20° and TR = 18ms. From images (i) and (iii) we computed a T1 map, from images (i) and (ii) we computed an MTR map, and from images (i), (ii) and (iii) we computed an MTsat map.

### Data acquisition

Three healthy male participants volunteered for multi-center data collection. Written informed consent was acquired prior to the data collection following a protocol approved by the Ethics Committee of each imaging center.

The participants (P1-3) and the ISMRM-NIST system phantom (HPD Inc., serial number = 42) were scanned on three imaging systems at two imaging sites. In S1 and S2, the phantom was scanned using a 20-channel head coil due to space constraints, whereas a 32-channel coil was used in G1. For in-vivo imaging, 32-channel head coils were used in G1 and S1, whereas a 64-channel coil was used in S2. S1 was equipped with an XR-K2309_2250V_951A (Siemens Healthineers, Erlangen,Germany) gradient system (80 mT/m maximum amplitude and 200 T/m/s slew rate per axis, 50 cm maximum FOV), S2 with an XQ-K2309_2250V_793A (Siemens Healthineers, Erlangen,Germany) gradient system (45 mT/m maximum amplitude and 200 T/m/s slew rate per axis, 50 cm maximum FOV) and G1 with a 8920-XGD (GE Healthcare, Milwaukee, USA) gradient system (50 mT/m maximum amplitude and 200 T/m/s slew rate per axis, 48cm maximum FOV). The nominal field strengths on G1 and S1-2 were 3T and 2.89T, respectively. Before the scan, the phantom was kept in the imaging site for at least a day, and in the scanner room for at least 3 hours. The measured bore temperature in G1, S1 and S2 was 20.1°C, 20.2°C and 20.8°C, respectively.

The acquisition parameters were set according to a generic protocol established for MTsat imaging of neural tissue^45^. The vendor-neutral acquisition parameters were identical on all systems. However, it was not possible to equalize all the parameters between the vendor-native protocols. Comparison of vendor-native and vendor-neutral protocols are presented in Table 1. To scan the phantom, prescan measurements were performed as described by Keenan et al. (2021) and the vendor-neutral acquisitions were configured to start the acquisitions with these calibrations^13^. For all acquisitions, the prescan settings of the initial T1w acquisition were used for the subsequent PDw and MTw acquisitions on all vendor systems. For the VENUS acquisitions, B0 shimming gradients were set using a spiral multi-echo gradient-echo sequence. Gradient non-linearity correction was performed as part of the on-site reconstruction pipeline. The warping coefficients were made available for offline reconstruction. For the systems S1-2, the identical protocol was used by exporting the vendor-native protocol files from S2. The protocols for G1 were set on-site.

**Table 1.**
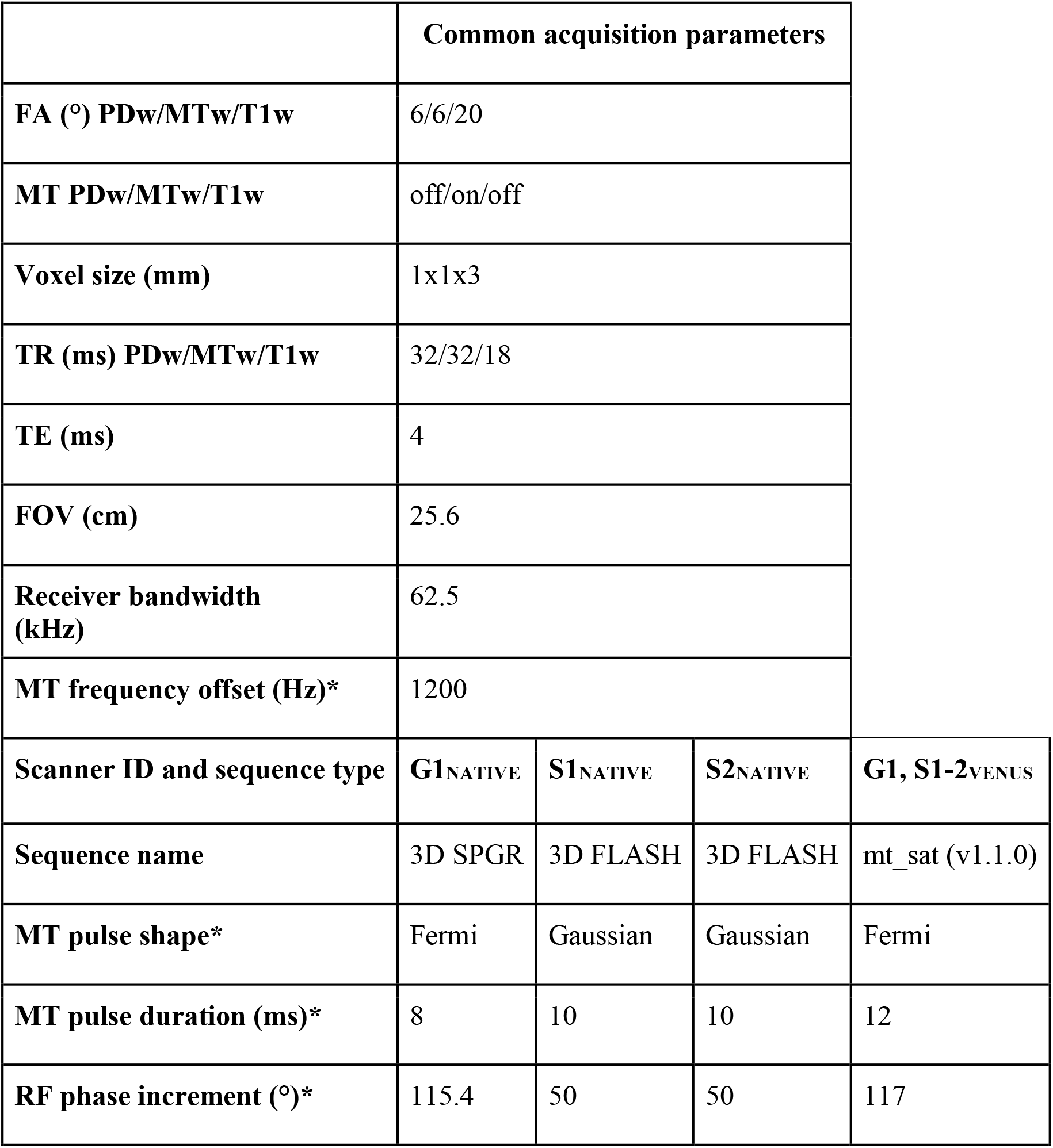
Comparison of acquisition parameters between vendor-native and vendor-neutral protocols. Parameters that are hardcoded on vendor-native systems are denoted by an asterisk (*).

|  | Common acquisition parameters |  |  |  |
| --- | --- | --- | --- | --- |
| **FA (°) PDw/MTw/T1w** | 6/6/20 |  |  |  |
| **MT PDw/MTw/T1w** | off/on/off |  |  |  |
| **Voxel size (mm)** | 1x1x3 |  |  |  |
| **TR (ms) PDw/MTw/T1w** | 32/32/18 |  |  |  |
| **TE (ms)** | 4 |  |  |  |
| **FOV (cm)** | 25.6 |  |  |  |
| **Receiver bandwidth (kHz)** | 62.5 |  |  |  |
| **MT frequency offset (Hz)\*** | 1200 |  |  |  |
| **Scanner ID and sequence type** | **G1<sub>NATIVE</sub>** | **S1<sub>NATIVE</sub>** | **S2<sub>NATIVE</sub>** | **G1, S1-2<sub>VENUS</sub>** |
| **Sequence name** | 3D SPGR | 3D FLASH | 3D FLASH | mt_sat (v1.1.0) |
| **MT pulse shape\*** | Fermi | Gaussian | Gaussian | Fermi |
| **MT pulse duration (ms)\*** | 8 | 10 | 10 | 12 |
| **RF phase increment (°)\*** | 115.4 | 50 | 50 | 117 |

### Data processing

All the processing was performed using data-driven and container mediated pipelines comprised of two docker images (Fig. 2). Quantitative fitting was performed in qMRLab^46^ v2.5.0b. Pre-processing steps were performed using ANTs^47^ for registration and FSL^48^ for automatic gray-matter (GM) and white-matter (WM) segmentation.

**Figure 2.**
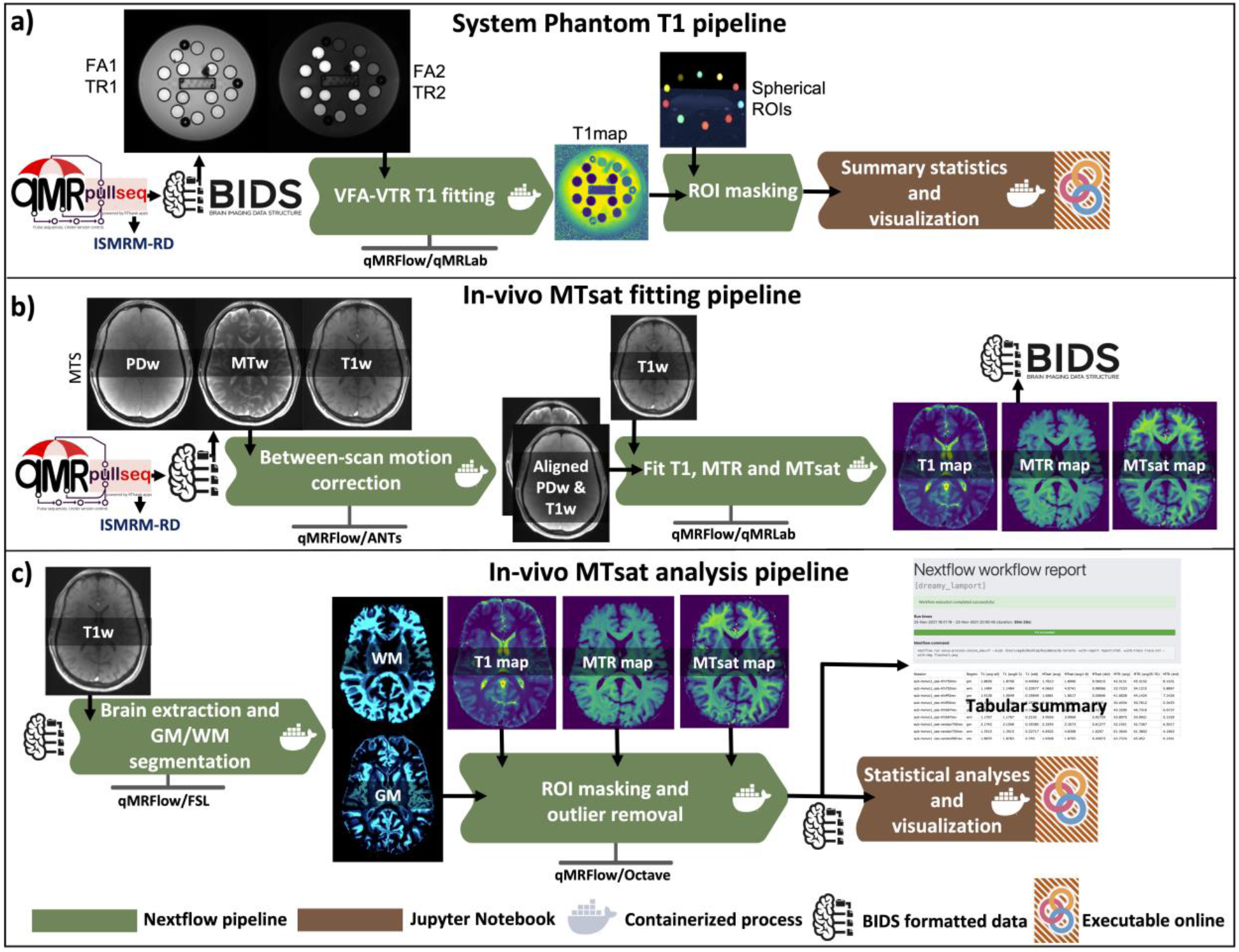
Image quality assessment using the phantom: **a)** Peak SNR values (PSNR) from T1w and PDw phantom images are displayed for vendor-neutral (red, orange, and yellow) and vendor-native (blue, cyan, and teal) G1, S1 and S2 scans, respectively. The same color coding is used in the following panels. **b-g)** Coronal PDw phantom images, with an inset zoom on two 4×4 grids with 1mm spacing. The brightness of the zoomed-in insets is increased by 30% for display purposes. **h-m**) Coronal T1w phantom images showing the center of the reference T1 arrays. The fine resolution (<0.6mm) inserts located at the center of the T1 array (rectangular area) are not relevant for the present resolution level. These inserts are colored following the same convention described in a) for convenience.

For the in-vivo data, between-scan motion correction was performed by aligning PDw and MTw images onto the T1w, followed by MTsat fitting (Fig. 2b). Brain region segmentations were performed on the T1w images and ROI masking was performed to prepare data for statistical analyses (Fig. 2c). The phantom T1 pipeline consisted of linearized VFA-T1 by accounting for varying TRs^22^ (Fig. 2a). Resultant phantom maps were then masked using spherical ROIs as described in^49^. Finally, peak SNR (PSNR) values were calculated, and the phantom images were visualized to compare image quality characteristics between vendor-native and VENUS implementations (Fig. 3).

**Figure 3.**
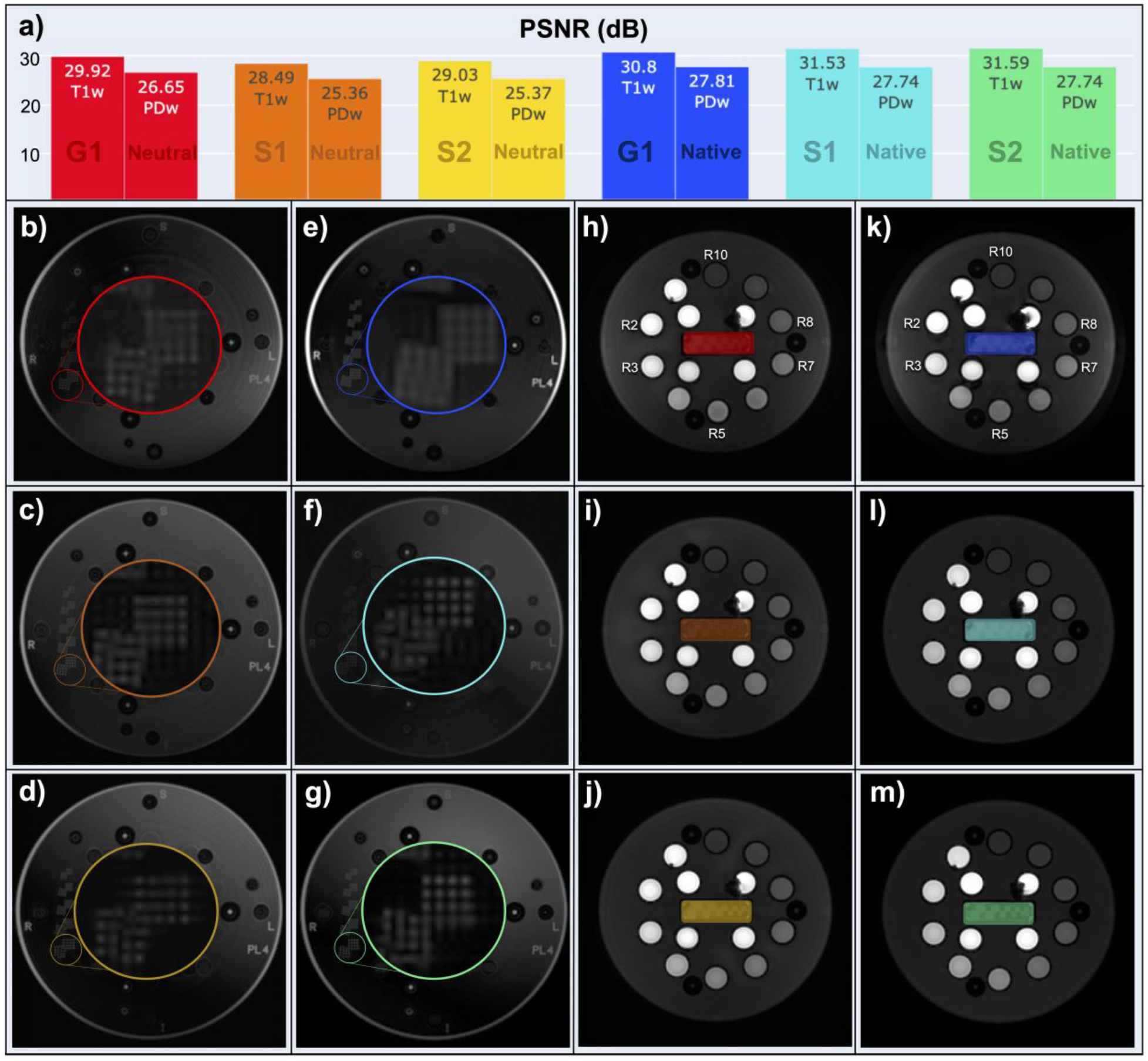
Overview of the analysis workflow for phantom scans (a) and in vivo scans (b, c). File collection (MTS) and output map names (T1map, MTsat, MTRmap) follow the BIDS standard v1.6.0. **a)** Vendor-neutral and vendor-native phantom images were acquired at two flip angles and two repetition times. The output data are then subjected to T1 fitting using qMRLab (Docker container image: qmrlab/minimal:v2.5.0b). The resulting T1 maps are masked using manually prescribed 10 spherical ROIs (reference T1 ranging from 0.9 to 1.9s). **b)** PDw and MTw images are aligned to the T1w image to correct for between-scan motion. The aligned dataset is then subjected to MTsat and MTR fitting in qMRLab to generate T1map, MTRmap and MTsat. **c)** Brain extraction and tissue type segmentation is performed on the T1w images using FSL. Following region masking and outlier removal for each map, vector outputs are saved for statistical analysis and visualization in an online-executable Jupyter Notebook (R-Studio and Python) environment. The tabular summary and the Nextflow pipeline execution report are exported. The pipeline execution report is available at https://qmrlab.org/VENUS/qmrflow-exec-report.html.

### Statistical analyses

All the descriptive statistics were reported by the processing pipeline in tabular format for phantom and in-vivo maps (available at https://osf.io/5n3cu). Vendor-neutral and vendor-native phantom measurement performances were compared against the reference (Fig. 4b,c) and percent deviations from the ground truth were reported (Fig. 4d).

**Figure 4.**
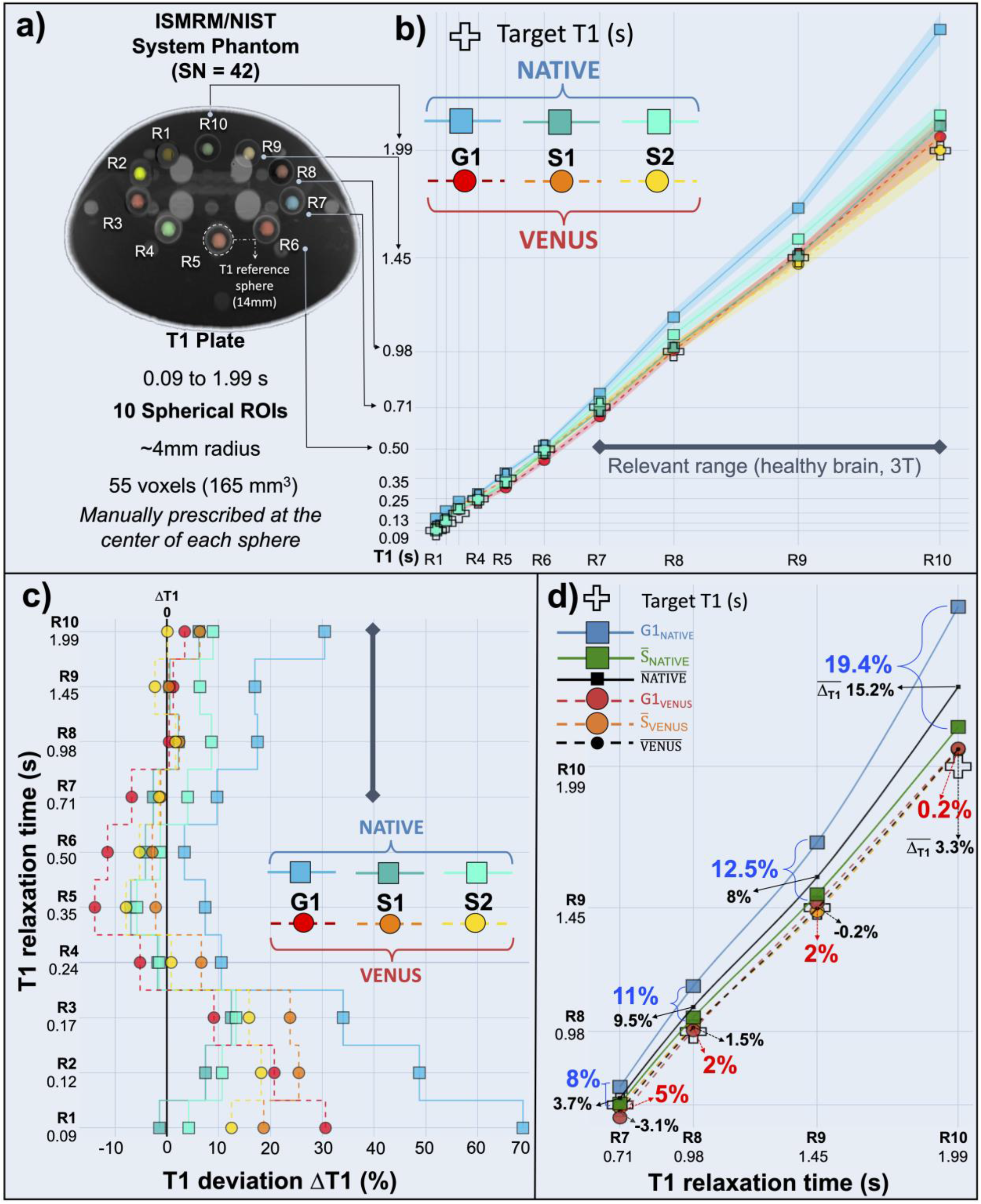
Comparison of vendor-native and vendor-neutral T1 measurements in the studied range of the phantom reference values, from 0.09 to 1.99s **(a)**. T1 values from the vendor-native acquisitions are represented by solid lines and square markers in cold colors, and those from VENUS attain dashed lines and circle markers in hot colors. **b)** Vendor-native measurements, especially G1_NATIVE_ and S2_NATIVE_, overestimate T1. G1_VENUS_ and S1-2_VENUS_ remain closer to the reference. **c)** For VENUS, ΔT1 remains low for R7 to R10, whereas deviations reach up to 30.4% for vendor-native measurements. **d)** T1 values are averaged over S1-2 (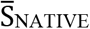 and 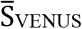, green square and orange circle) and according to the acquisition type (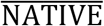 and 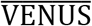, black square and black circle). Inter-vendor percent differences are annotated in blue (native) and red (VENUS). Averaged percent measurement errors 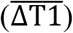 are annotated on the plot (black arrows).

Kernel density estimates of the T1, MTR and MTsat distributions in WM and GM were visualized as ridgeline plots for one participant (Fig. 5d-i). Before the statistical comparisons in WM, the outliers were removed from the distributions. The non-outlier range was 0 to 3s for T1, 35 to 70% for MTR and 1 to 8 for MTsat. Filtered distributions were then randomly sampled to obtain an equal number of WM voxels (N = 37,000) for a balanced comparison.

**Figure 5.**
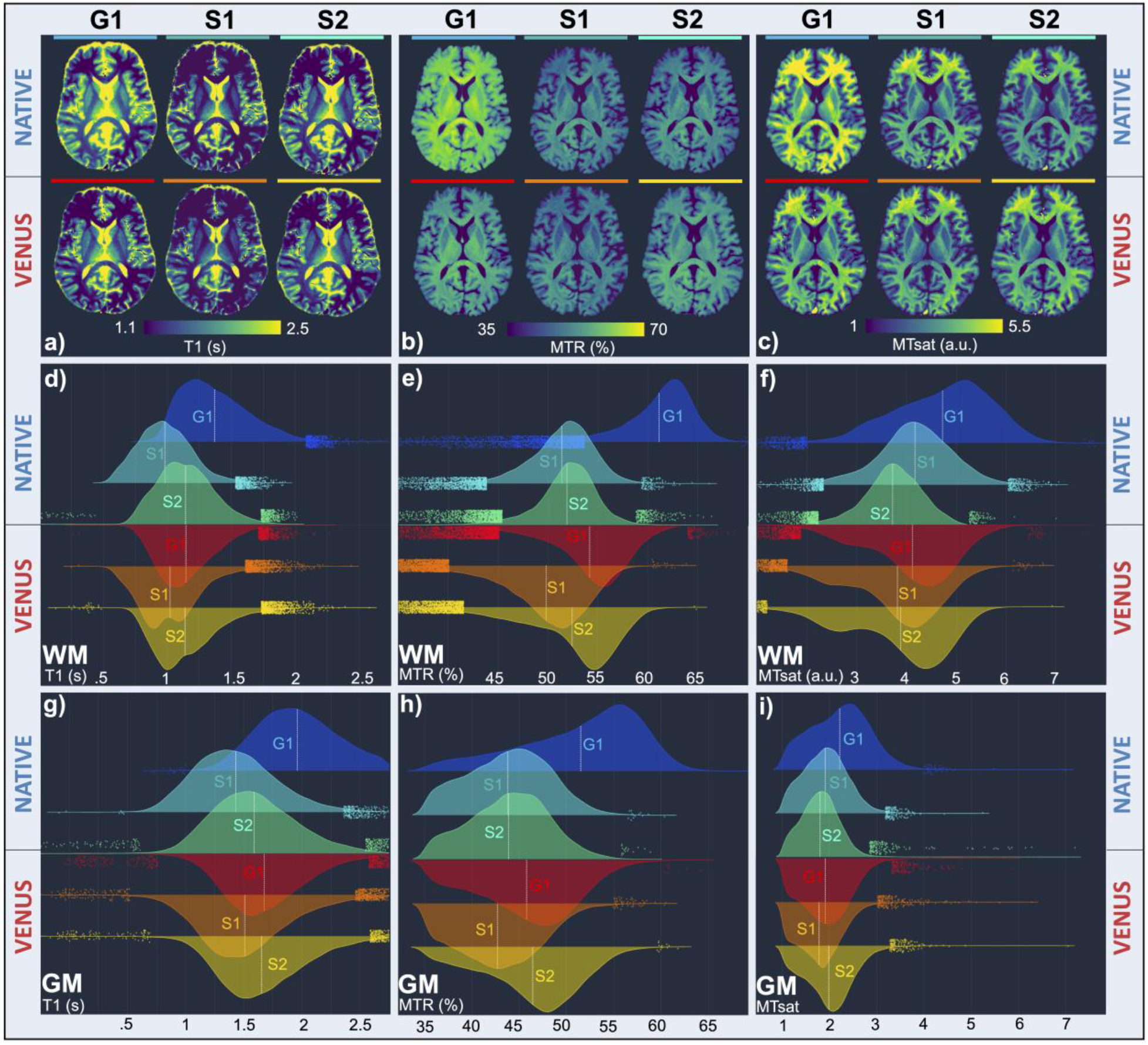
Vendor-native and VENUS quantitative maps from one participant are shown in one axial slice (a-c). Distributions of quantified parameters in white matter (d-f) and gray matter (g-i) are shown using ridgeline plots of kernel density estimations. **a-c)** Inter-vendor images (G1 vs S1 and G1 vs S2) appear more similar in VENUS (lower row) than in native (upper row). **d-f)** Distribution shapes and locations agree with visual inspection from (a), indicating closer agreement between VENUS distributions. **g-i)** Superior between-scanner agreement of VENUS persists in GM as well. Compared to WM, GM distributions are in the expected range (higher T1, lower MTR and MTsat values).

Percentile bootstrap based shift function analysis^50^ was performed to compare dependent measurements of T1, MTR and MTsat in WM (N=37,000) between different systems (G1-vs-S1, G1-vs-S2 and S1-vs-S2) for VENUS and for the vendor-native implementations. Deciles of the distributions were computed using a Harrell-Davis quantile estimator^51^. The decile differences were calculated using 250 bootstrap samples to characterize differences at any location of the distributions (Fig. 6a). For convenience, we annotated the 5th decile (median) with the respective percent difference (Fig. 6b-d). To characterize the difference between scanners across the participants, the shift function was extended to a hierarchical design (Fig. 7a). Similarly, percent T1, MTR and MTsat differences between scanners at the median deciles were annotated per subject, and the average percent deviations were reported (Fig. 7b-d). The reader is welcome to reproduce these figures online, where necessary changes can be made to visualize the high-density intervals of the decile differences at https://github.com/qMRLab/VENUS.

**Figure 6.**
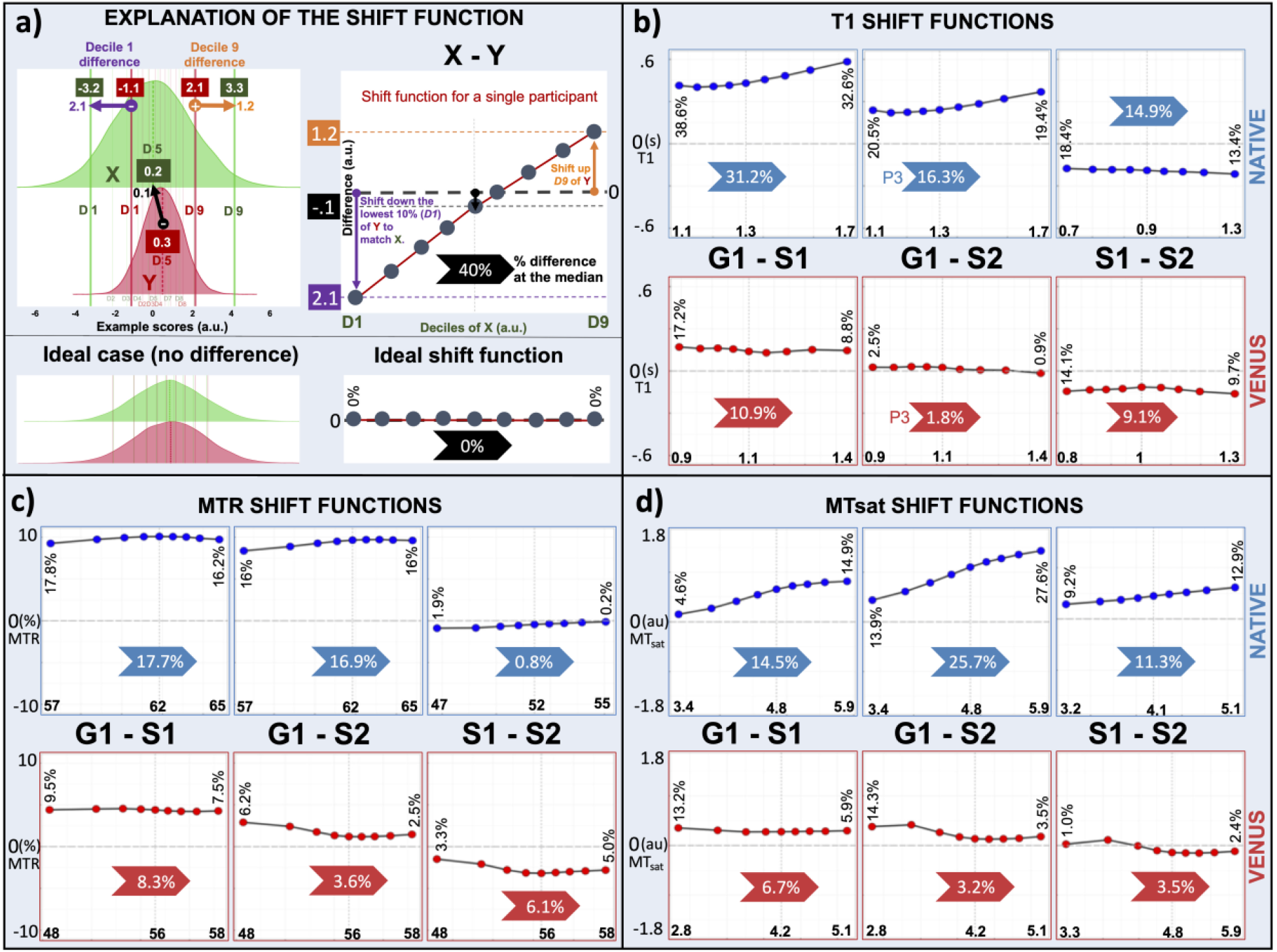
Shift function analysis of T1, MTR and MTsat results from a single participant in white-matter (WM). **a)** Shift function analysis is a graphical tool for analyzing differences between two (dependent in this case) measurements at any location of the distributions. It shows 9 markers dividing the distribution into 10 equal chunks; hence the markers represent deciles. The shape of the curve (shift function) obtained by plotting decile differences against the first decile characterizes how distributions differ from each other. **b-d)** Here, shift function plots compare the agreement between different scanners for VENUS (bottom row) and vendor-native (top row) implementations in quantifying T1, MTR and MTsat. Across all the comparisons, the apparent trend is that the VENUS inter-vendor variability is lower than for the vendor-native implementations.

**Figure 7.**
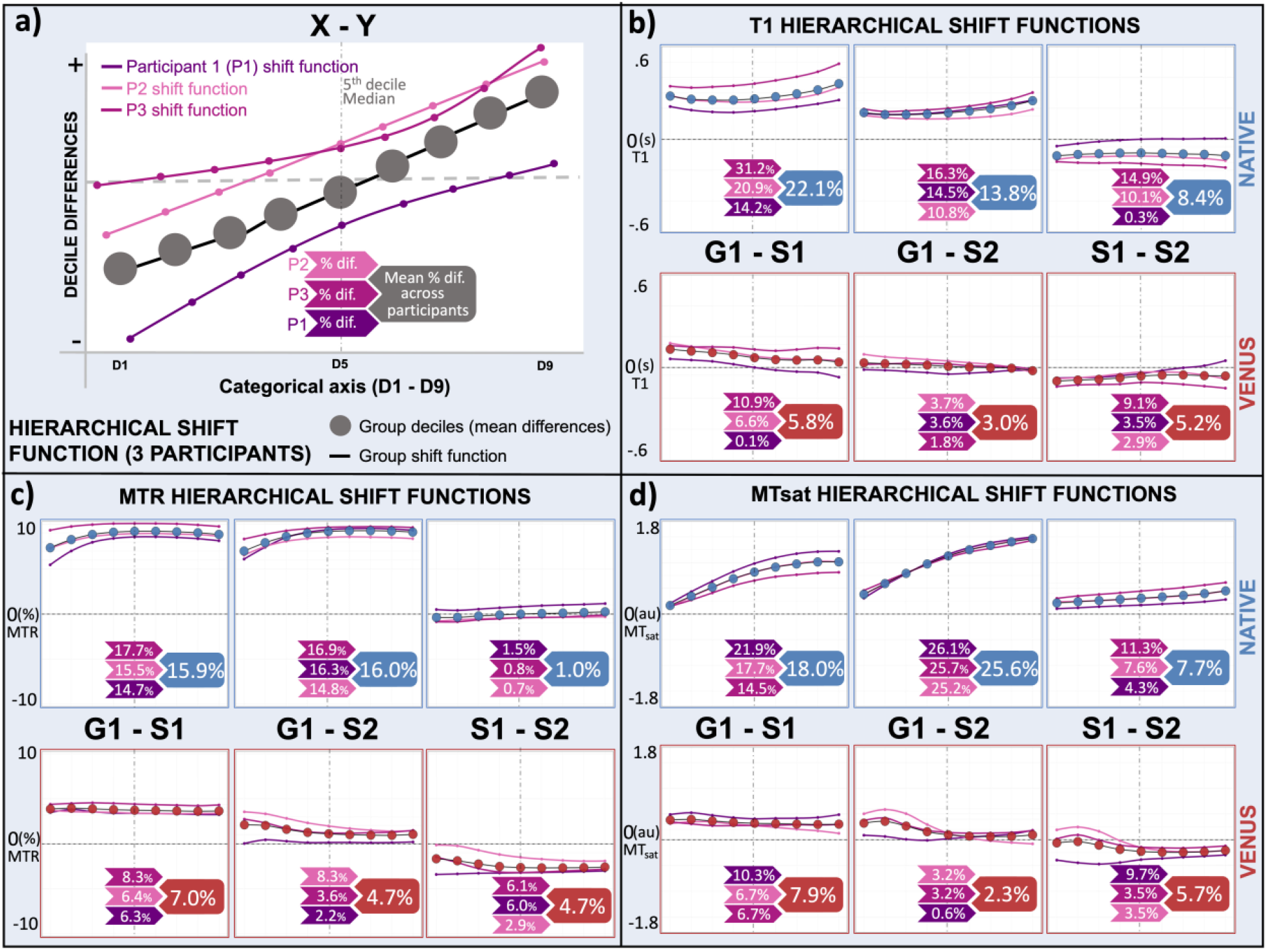
Hierarchical shift function analysis of T1, MTR and MTsat results from three participants in the white-matter (WM). **a)** Hierarchical shift function repeats Figure 6 for all participants (shades of pink). Group deciles (red and blue markers for VENUS and vendor-native, respectively) show the average trend of inter-scanner differences across participants. **b-d)** G1-vs-S1 and G1-vs-S2 (inter-vendor) agree in VENUS better than they do in vendor-native for all quantitative maps of T1, MTR and MTsat.

Finally, quantitative measurement discrepancies of vendor-native and VENUS implementations between different vendors were compared using Wilcoxon signed rank test. The comparison was performed on the G1-vs-S1 and G1-vs-S2 percent absolute differences of T1, MTR and MTsat in white matter between vendor-native and vendor-neutral implementations. The level of significance was set at p = 0.05.

## Results

The contrast characteristics of VENUS and vendor-native T1w phantom images are qualitatively comparable (Fig. 3h-m). In addition, VENUS PSNR values are on a par with those of vendor-native T1w and PDw images (Fig. 3a). The resolution markers are discernible in the vendor-neutral images (Fig. 3b-d) with a slightly lower horizontal resolution compared to the S1-2_NATIVE_ (Fig. 3f,g). On the other hand, the insert pattern resolution of G1_NATIVE_ (Fig. 3e) appears lower than that of G1_VENUS_ (Fig. 3b).

Overall, the vendor-neutral implementation reduces inter-vendor variability and brings T1 estimations closer to the ground truth of the phantom, particularly for the targeted physiological interval from 0.7 to 1.9s (Fig. 4b). On the other hand, T1 deviations (ΔT1) calculated by percent error indicate that G1_NATIVE_ and S2_NATIVE_ exhibit a persistent overestimation trend, with S1_NATIVE_ showing a relatively better accuracy (Fig. 4c). Within the same interval, the highest deviation is observed for G1_NATIVE_, where ΔT1 ranges from 9.7 to 30.4%. For R4-6, G1_NATIVE_ and G1_VENUS_ T1 measurements straddle the reference, where G1_VENUS_ shows 5.1-13.8% underestimation and the G1_NATIVE_ overestimation remains within the 3.4-10.5% interval (Fig. 4c). For lower T1 reference values (T1 < 170ms), all measurements indicate higher deviations, with S1-2_NATIVE_ performing better than S1-2_VENUS_.

When the measured T1 values are averaged over S1-2 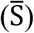, the differences between G1_NATIVE_ and 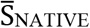 are 8, 11, 12.5 and 19.4%, whereas the differences between G1_VENUS_ and 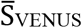 are 5, 2, 2 and 0.2% for R7-10, respectively (Fig. 4d). This reduction in between-vendor differences brought by VENUS is coupled with an improvement in accuracy. When averaged according to the implementation type, average VENUS deviation 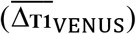 falls within the 0.2 - 4% range and 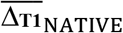 ranges from 7 to 11%. Even though G1_NATIVE_ has the dominant contribution to the higher 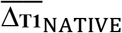 values, Fig. 4d shows that 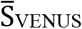 is closer to the reference than 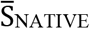 for most of the R7-10 (ΔT1 of 7.6, 3.5, 5.4, 0.7% and 3.2, 0.9, 2, 1.3% for 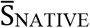 and 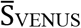, respectively). As a result, VENUS reduces between-vendor differences with an overall accuracy improvement. Figure 5 shows in vivo T1, MTR and MTsat maps from a single participant (P3). While most of the improvements are evident from the maps (5a - 5c), the ridgeline plots (5d – 5i) make it easier to appreciate the VENUS vs vendor-native distribution differences in the GM and WM per metric.

Consistent with the higher myelin content in WM, T1 values are lower in WM (around 1.1 ∓ 0.2s, Fig. 5d), whereas MTR and MTsat values are higher (around 50 ∓ 8% and 3.8 ∓ 0.9 a.u., Fig. 5 e,f) in comparison to those in GM (1.9 ∓ 0.4s, 40 ∓ 2% and 1.8 ∓ 0.5, for T1, MTR and MTsat, respectively, Fig. 5g-i). The general trend observed in the images is captured by ridgeline plots, showing better agreement between VENUS distributions of G1, S1 and S2. This is further supported by the between-scanner coefficient of variation (CoV) per metric (Table 2), showing that VENUS reduces the CoV from 16.5, 10.1 and 12.5% to 6.1, 4.1 and 4.1% for T1, MTR and MTsat, respectively. This indicates a sizable decrease in between-scanner variability using VENUS compared with vendor-native measurements and the trend is consistent across participants.

**Table 2.** Coefficient of variation (%) of vendor-neutral (VENUS) and vendor-native quantitative measurements between the scanners for each participant (P1-P3) and across participants.

| **Participants** | **P1** |  | **P2** |  | **P3** |  | **Across** |  |
| --- | --- | --- | --- | --- | --- | --- | --- | --- |
| **Protocol** | **NATIVE** | **VENUS** | **NATIVE** | **VENUS** | **NATIVE** | **VENUS** | **NATIVE** | **VENUS** |
| **T1** | 9.8 | 1.3 | 11.7 | 4.0 | 16.5 | 6.1 | 11.36 | 4.3 |
| **MTR** | 8.5 | 3.4 | 8.5 | 3.4 | 10.1 | 4.1 | 7.9 | 3.2 |
| **MTsat** | 13.6 | 5.9 | 11.9 | 3.4 | 12.1 | 4.1 | 10.7 | 4.2 |

Going from vendor-native (top rows, blue panels) to VENUS (bottom rows, red panels), Fig. 6b-d indicates a decrease in T1, MTR and MTsat WM differences between scanners from different vendors (G1-vs-S1 and G1-vs-S2) for P3, without exception and throughout the deciles. One can also appreciate the changes in shift function shapes. For example, the shift function for G1NATIVE vs S2_NATIVE_ MTsat comparison in Fig. 6d shows a positive linear trend, indicating that WM voxels with higher MTsat values tend to show a higher between-vendor difference. On the other hand, the G1_VENUS_ vs S2_VENUS_ MTsat shift function appears flatter, describing a more uniform (and reduced) bias throughout the WM distribution. As for within-vendor comparisons (S1-vs-S2) of the same participant, VENUS reduces difference scores for T1 and MTsat by 5.8 and 7.8% while increasing that for MTR by 5.3% (Fig. 6).

Figure 7 expands on Figure 6 for multiple participants by overlaying individual shift functions (shades of pink) and illustrating the across-participants trend using group shift functions that are red for VENUS and blue for vendor-native differences (Fig. 7a). Overall, VENUS G1-vs-S1 and G1-vs-S2 differences are on the order of 2.3 to 7.9%, whereas the vendor-native variations start from 13.8% and extends up to 25.6%, averaged across participants. The reduction in between-vendor differences achieved by VENUS is significant after correction for multiple comparisons for all maps (p=0.015). Another general observation is that individual shift function shapes are mostly consistent across participants, indicating that the inter-scanner differences between the VENUS and vendor-native implementations are not modified by anatomical differences. However, the magnitude of the difference is participant-specific. For example, P3 shows the highest G1_NATIVE_ vs S1_NATIVE_ T1 difference of 31.2%, which is followed by 20.9 and 14.2% for P2 and P1, respectively (Fig. 7b). Finally, within-vendor effect of VENUS remains on the lower side with all participants considered, reducing the S1-vs-S2 difference by 3.2 and 2.0% for T1 and MTsat while increasing that for MTR by 3.7%.

## Discussion

In this study, we developed and deployed a vendor-neutral qMRI protocol (VENUS) for T1, MTR and MTsat mapping on three 3T commercial scanners by two vendors. Our findings confirm the hypothesis that vendor-neutral sequences decrease inter-vendor variability of T1, MTR and MTsat measurements. This key improvement addresses *problem 1*, as stated in the Introduction, with open-source pulse sequence descriptions. The developed sequence can be run on most GE and Siemens scanners through RTHawk software and an additional UUI that allows users to prescribe customized file naming entities for exporting reconstructed images in the BIDS^43,44^ and k-space data in the ISMRM-RD format^42^. Conforming with community data standards, providing a user-friendly solution with a simplified vendor-neutral deployment, this work offers a complete solution for *problem 2,* and shows a way forward for the standardization of qMRI.

### Developing an end-to-end qMRI workflow

First, we created a vendor-native qMRI protocol that is unified across vendors to the greatest extent possible, by keeping contrast, timing, and acquisition geometry identical (Table 1). However, other vendor-native implementation details such as RF spoiling, MT and excitation pulse characteristics were different, as it is commonly the case in multicenter studies^14,38^. Trying to address these issues is difficult with a vendor-native sequence given that the implementations of commercial stock sequences commonly used for qMRI are not open *(problem 1).* One candidate solution for this problem is modifying sequences on the vendor’s proprietary development environment to equalize implementations as much as possible, which has been shown to improve reproducibility to some extent^21^. However, this requires familiarity with multiple sequence development environments and still may fall short in unifying all the aspects on-site. Not only is this approach impractical for the developers, but it is also not a user-friendly solution for clinical use. As we mention in the context of *problem 2,* reproducibility solutions unifying inter-vendor implementations become more favorable if they are designed with clinicians’ needs in mind. To that end, we aimed at providing a unified and smooth user experience by developing VENUS as an RTHawk application, which allows implementation details to be shared publicly starting at the pulse sequence level.

Second, we built from scratch a vendor-neutral sequence that was developed and tested on a single site and then ported to two more scanners from different vendors. In doing so, we adapted a system that is primarily geared toward real-time imaging (RTHawk) to perform quantitative MRI measurements. For example, absolute gradient limits have been allowed to achieve higher spoiling gradient moments and string-valued customized metadata injection has been enabled to follow community data standards^42–44^.

Third, we created a fully transparent, container-mediated and data-driven workflow^52,53^ that automates the processing and reduces variability introduced by the operators. By design, the workflow operates according to the BIDS qMRI standard^44^ for picking up all the necessary data and metadata, and generates outputs following a consistent derivative hierarchy. Moreover, the raw data is exported in the ISMRM-RD format by our vendor-neutral sequence, allowing the use of community developed reconstruction tools by simply adding another container at the beginning of our modular workflow. We envision that using open-source reconstruction tools would be highly favourable for vendor-neutral sequences employing under-sampled k-space with complex trajectories to guarantee reproducibility^54–57^.

### Reducing inter-vendor variability

Stock sequences are optimized for reliable clinical imaging. These optimizations do not necessarily serve for accuracy when the sequences are used for qMRI experiments. For example, the phase increment values of S1-2_NATIVE_ sequences (Table 1) are hardcoded to maximize in-vivo signal stability^58^, not T1 accuracy in phantoms^59^. On the other hand, the phase increment of G1 has been shown to be unsuitable for T1 mapping, exhibiting severe overestimations^14^. In this study, we set the value (117°) suggested for T1 accuracy^14^ while unifying all other aspects of the vendor-neutral acquisition between scanners. The results from the phantom analysis clearly demonstrate that VENUS achieves higher accuracy and a notable reduction in inter-vendor variability compared to its vendor-native counterparts (Fig. 4).

In the absence of an in-vivo ground-truth T1 map (from inversion recovery), we only looked at the agreement between the three implementations and explored whether VENUS brought the T1 values closer across vendors when compared to the vendor-native sequences. Visually (Fig. 5a), the reduction in T1 variability can be appreciated for VENUS within the dynamic range of T1 adjusted for WM/GM. As supported by the ridgeline plots (Fig. 5d,g), the G1_NATIVE_ T1 distribution is globally shifted towards higher values compared to S1-2_NATIVE_, and their central tendency differs. As observed in the phantom, G1_VENUS_ alleviates this discrepancy, shifting the T1 distribution closer to those of S1-2_VENUS_. Interestingly, the WM T1 distributions appear more unimodal on G1 compared to S1-2 (both for VENUS and vendor-native), with a more pronounced bimodal appearance for S1-2_VENUS_. A plausible explanation for that are vendor and implementation specific differences due to B1+ field inhomogeneity. Nevertheless, the VENUS shift functions for G1-vs-S1 and G1-vs-S2 comparisons are flatter than the vendor-native shift functions (Fig. 6b), indicating that the inter-vendor WM T1 statistical distribution characteristics are more similar using VENUS.

Table 2 indicates that reduction in inter-vendor variability is not limited to T1 but persists for all the metrics across all participants. The inter-vendor variability in MTR and MTsat is relatively easier to appreciate visually (Fig. 5b,c). The three MTR and MTsat maps from VENUS are in better agreement, and this is most likely because our unified implementation compensated for the MT saturation pulse differences (Table 1).

Reducing variability matters as much as which tools we use to assess it. Shift functions^50^ take the comparison beyond differences in point estimates of centrality and relative spread (CoV) to a robust characterization of differences on the absolute scale of the measurement. This makes the shift function analysis (Fig. 6-7) more informative than CoV (Table 2) by characterizing how distributions differ for P3. For example, Table 2 shows that VENUS reduces CoV from 12.1 to 4.1% for MTsat. Figure 6d explains that most of that reduction is achieved by decreasing the absolute G1-vs-S2 MTsat difference from 1.1 to 0.1 (a.u.), corresponding to a reduction of the inter-vendor difference from 25.7% to 3.2%. In addition, Fig. 6d indicates that higher deciles benefit from the G1-vs-S2 variability reduction more compared to the lower deciles, yielding a flatter shift function for VENUS. This suggests that VENUS not only brings averaged MTsat values closer, but also matches their distribution shape (Fig. 5f).

### Implications of vendor-neutrality and the importance of transparency

The most important contribution of this article is the vendor-neutral solution it provides for multi-center qMRI by significantly reducing inter-vendor variability. This issue has been hampering the standardization of qMRI methods for multi-center clinical trials^60^, validation^61,62^, establishing protocols^17^, applied neuroimaging studies^63^, determining the range of parameters in pathology^64,65^ and in health^16,38^, scanner upgrades^49^ and even for phantom studies^12,13^. By reducing such variabilities, the VENUS approach can bring qMRI closer to teasing out the true biological variability in quantifying in-vivo tissue microstructure^66^.

We recognize that part of the RTHawk workflow is proprietary. Hence, we emphasize the importance of the transparency to inter-vendor reproducibility at the level of sequence definitions. RTHawk allows sharing open-source sequences (https://github.com/qMRLab/mt_sat). Note that neither RTHawk nor open-source solutions can access under the hood of vendor-specific drivers to guarantee that open-source sequences are executed according to the published code. To achieve such open-execution^67^, vendor-neutral solutions should be coupled with open-hardware^68^. Although RTHawk’s pulse sequence and data management servers give more flexibility to the scanner operation at multiple levels of the workflow (e.g., UUI, customized raw data stream, asynchronous real-time updates to sequences, standalone workstation etc.), the conversion of the open-source sequence descriptions to vendor-specific hardware instructions is not transparent. We argue that this is a reasonable trade-off as it peels another layer from a vendor-specific ecosystem, and it does not sacrifice the transparency of sources relevant to a pulse sequence description. The accuracy and reliability of the parameter estimation depend on these descriptions; therefore, for qMRI to work we need to be able to access, modify, and share the methods^69^. Fortunately, the VENUS approach to qMRI is not framework-exclusive and satisfies this key requirement.

Namely, using community developed tools such as Pulseq, GammaStar, SequenceTree, ODIN or TOPPE, interoperable qMRI applications can be developed. A critical step to achieve this is effective communication between method developers to foster compatibility between frameworks. This is nicely exemplified by GammaStar and JEMRIS, as both applications can export Pulseq descriptions. Enabling a similar feature by developing a SpinBench plugin is among our future goals. To facilitate discussions on this topic with vendor-neutral framework developers, we created a forum page on the code repository of this article (https://github.com/qMRLab/VENUS).

### Limitations and future directions

The RF transmission systems were different between all the scanners used for data collection. This is indeed a likely cause of variability of T1 and MTsat maps. Therefore, another obvious limitation of this study is the lack of B1+ mapping. Unfortunately, a vendor-native B1+ mapping sequence was not available on G1, and it is also well-known that discrepancies between vendor-native B1+ mapping contribute to between-scanner bias in T1 mapping^38^. As for the VENUS protocol, the current version of RTHawk did not permit the long gradient durations (e.g., 80ms) needed by AFI implementation to achieve accurate B1+ mapping^70^. Therefore, further investigation is needed to compare vendor-neutral B1+ maps across vendors for isolating the specific contribution of transmit field inhomogeneity.

Another critical factor affecting the accuracy is the calculation of a global RF scaling factor. Vendor-native systems set the transmit gain using their own prescan routine, which may lead to a systematic bias in quantitative mapping. In this work, we implemented prescan for G1 and S1-2 as described by^13^ and configured RTHawk to use the same calibration measurements. Nevertheless, it is possible to make this step vendor-neutral as well. For future work, we plan to develop a double-angle VENUS prescan using the same excitation pulses as the qMRI sequences that follow, to determine a global RF scaling factor. Coupled with the use of anatomy-mimicking quantitative MRI phantoms^71^, this would offer qMRI-ready adaptive prescan routines and help investigate the effect of standardizing calibration measurements on multicenter accuracy and agreement.

We made the details of the RTHawk reconstruction pipeline publicly available. However, the raw data from the vendor-native acquisitions were not available. Open-source reconstruction tools^54–56^ are an important asset to investigate the potential effect of reconstruction pipeline differences on image characteristics, such as the differences between resolution insert patterns observed in Fig. 3b-g. Therefore, future work will enable raw data export from vendor-native systems and add a containerized reconstruction node to the qMRFLow^53^ pipeline for investigating potential sources of reconstruction variability.

Finally, the study of measurement stability using VENUS could benefit from recruiting more participants and including more imaging sites. Although the inter-vendor pattern observed in our limited cohort is consistent across 3 scanners, within-vendor (S1-vs-S2) results from vendor-native implementations are more consistent and comparable to VENUS (Fig. 7). Nevertheless, more data is needed for a thorough characterization of subject specific within-vendor effects. Our future study will deploy VENUS on more GE and Siemens sites and recruit more participants to investigate the variability problem from different perspectives, including system upgrades and WM pathology.

## Conclusion

In this article we have demonstrated that vendor-neutral sequences and transparent workflows reduce inter-vendor variability in quantitative MRI. Additionally, these workflows can be deployed on an FDA-approved device, which demonstrates the potential for wide clinical adoption. Quantitative MRI needs to bypass the vendor black boxes to make an impact in the clinic, and this work shows the way forward.

## Acknowledgement

The authors would like to acknowledge Graham Wright, PhD for his help in organizing the multicenter experiment, Paule Samson for her help in data collection, Juan Santos, PhD and William R. Overall, PhD for their technical support in deploying RTHawk on multiple sites.

## Data availability statement

All the vendor-neutral pulse sequences are publicly available as git submodules at https://github.com/qmrlab/pulse_sequences and can be run on RTHawk systems v3.0.0 and later. The RF and gradient waveforms (spv files) can be inspected and simulated using SpinBench (https://www.heartvista.ai/spinbench). As per the general design principles of fully reproducible qMRFlow pipelines, we adhered to a one-process one-container mapping for the processing of this dataset. Docker images, BIDS and ISMRM-RD compliant dataset from the current study are freely available at https://doi.org/10.17605/osf.io/5n3cu. Finally, the whole analysis and interactive version of all the figures in this article will be available and executable online at https://github.com/qmrlab/venus.

## References

1. Lauterbur PC. Image formation by induced local interactions: examples employing nuclear magnetic resonance. nature. 1973;242(5394):190–191. doi:10.1038/242190a0

2. Gillies RJ, Kinahan PE, Hricak H. Radiomics: Images Are More than Pictures, They Are Data. Radiology. Feb 2016;278(2):563–77. doi:10.1148/radiol.2015151169

3. Novikov DS, Kiselev VG, Jespersen SN. On modeling. Magn Reson Med. Jun 2018;79(6):3172–3193. doi:10.1002/mrm.27101

4. Hahn EL. An Accurate Nuclear Magnetic Resonance Method for Measuring Spin-Lattice Relaxation Times. Phys Rev. 1949;76(1):145–146. doi:10.1103/PhysRev.76.145

5. Gupta RK. A new look at the method of variable nutation angle for the measurement of spin-lattice relaxation times using fourier transform NMR. Journal of Magnetic Resonance. 1977;25(1):231–235. doi:10.1016/0022-2364(77)90138-X

6. D.C. Look, Locker DR. Time saving in measurement of NMR and EPR relaxation times. Review of Scientific Instruments. 1970;41(2):250–251. doi:10.1063/1.1684482

7. Marques JP, Kober T, Krueger G, van der Zwaag W, Van de Moortele PF, Gruetter R. MP2RAGE, a self bias-field corrected sequence for improved segmentation and T1-mapping at high field. Neuroimage. Jan 15 2010;49(2):1271–81. doi:10.1016/j.neuroimage.2009.10.002

8. Stikov N, Boudreau M, Levesque IR, Tardif CL, Barral JK, Pike GB. On the accuracy of T1 mapping: searching for common ground. Magn Reson Med. Feb 2015;73(2):514–22. doi:10.1002/mrm.25135

9. Grafe D, Frahm J, Merkenschlager A, Voit D, Hirsch FW. Quantitative T1 mapping of the normal brain from early infancy to adulthood. Pediatr Radiol. Mar 2021;51(3):450–456. doi:10.1007/s00247-020-04842-7

10. Okubo G, Okada T, Yamamoto A, et al. MP2RAGE for deep gray matter measurement of the brain: A comparative study with MPRAGE. J Magn Reson Imaging. Jan 2016;43(1):55–62. doi:10.1002/jmri.24960

11. Voelker MN, Kraff O, Goerke S, et al. The traveling heads 2.0: Multicenter reproducibility of quantitative imaging methods at 7 Tesla. Neuroimage. May 15 2021;232:117910. doi:10.1016/j.neuroimage.2021.117910

12. Bane O, Hectors SJ, Wagner M, et al. Accuracy, repeatability, and interplatform reproducibility of T 1 quantification methods used for DCE-MRI: Results from a multicenter phantom study. Magnetic Resonance in Medicine. 2018;79(5):2564–2575. doi:10.1002/mrm.26903

13. Keenan KE, Gimbutas Z, Dienstfrey A, et al. Multi-site, multi-platform comparison of MRI T1 measurement using the system phantom. PLOS ONE. 2021;16(6):e0252966. doi:10.1371/journal.pone.0252966

14. Yarnykh VL. Optimal radiofrequency and gradient spoiling for improved accuracy of T1 and B1 measurements using fast steady-state techniques. Magnetic Resonance in Medicine. 2010;63(6):1610–1626. doi:10.1002/mrm.22394

15. A.G. Teixeira RP, Malik SJ, Hajnal JV. Fast quantitative MRI using controlled saturation magnetization transfer. Magnetic Resonance in Medicine. 2019;81(2):907–920. doi:10.1002/mrm.27442

16. Bojorquez JZ, Bricq S, Acquitter C, Brunotte F, Walker PM, Lalande A. What are normal relaxation times of tissues at 3 T? Magn Reson Imaging. Jan 2017;35:69–80. doi:10.1016/j.mri.2016.08.021

17. Cohen-Adad J, Alonso-Ortiz E, Abramovic M, et al. Open-access quantitative MRI data of the spinal cord and reproducibility across participants, sites and manufacturers. Sci Data. Aug 16 2021;8(1):219. doi:10.1038/s41597-021-00941-8

18. Liberman G, Louzoun Y, Ben Bashat D. T(1) mapping using variable flip angle SPGR data with flip angle correction. J Magn Reson Imaging. Jul 2014;40(1):171–80. doi:10.1002/jmri.24373

19. Leutritz T, Seif M, Helms G, et al. Multiparameter mapping of relaxation (R1, R2*), proton density and magnetization transfer saturation at 3 T: A multicenter dual-vendor reproducibility and repeatability study. Hum Brain Mapp. Oct 15 2020;41(15):4232–4247. doi:10.1002/hbm.25122

20. Boudreau M, Stikov N, Pike GB. B1 -sensitivity analysis of quantitative magnetization transfer imaging. Magn Reson Med. Jan 2018;79(1):276–285. doi:10.1002/mrm.26673

21. Gracien R-M, Maiworm M, Brüche N, et al. How stable is quantitative MRI? – Assessment of intra-and inter-scanner-model reproducibility using identical acquisition sequences and data analysis programs. NeuroImage. 2020;207:116364. doi:10.1016/j.neuroimage.2019.116364

22. Helms G, Dathe H, Dechent P. Quantitative FLASH MRI at 3T using a rational approximation of the Ernst equation. Magn Reson Med. Mar 2008;59(3):667–72. doi:10.1002/mrm.21542

23. Rowley CD, Campbell JSW, Wu Z, et al. A model-based framework for correcting B 1 + inhomogeneity effects in magnetization transfer saturation and inhomogeneous magnetization transfer saturation maps. Magn Reson Med. Oct 2021;86(4):2192–2207. doi:10.1002/mrm.28831

24. Stikov N, Trzasko JD, Bernstein MA. Reproducibility and the future of MRI research. Magn Reson Med. Dec 2019;82(6):1981–1983. doi:10.1002/mrm.27939

25. Jochimsen TH, Von Mengershausen M. ODIN—object-oriented development interface for NMR. Journal of Magnetic Resonance. 2004;170(1):67–78. doi:10.1016/j.jmr.2004.05.021

26. Stöcker T, Vahedipour K, Pflugfelder D, Shah NJ. High-performance computing MRI simulations. Magnetic resonance in medicine. 2010;64(1):186–193. doi:10.1002/mrm.22406

27. Magland JF, Li C, Langham MC, Wehrli FW. Pulse sequence programming in a dynamic visual environment: SequenceTree. Magnetic resonance in medicine. 2016;75(1):257–265. doi:10.1002/mrm.25640

28. Ravi KS, Geethanath S, Vaughan JT. PyPulseq: A python package for mri pulse sequence design. Journal of Open Source Software. 2019;4(42):1725.

29. Layton KJ, Kroboth S, Jia F, et al. Pulseq: a rapid and hardware-independent pulse sequence prototyping framework. Magnetic resonance in medicine. 2017;77(4):1544–1552. doi:10.1002/mrm.26235

30. Nielsen JF, Noll DC. TOPPE: A framework for rapid prototyping of MR pulse sequences. Magnetic resonance in medicine. 2018;79(6):3128–3134. doi: 10.1002/mrm.26990

31. Cordes C, Konstandin S, Porter D, Günther M. Portable and platform-independent MR pulse sequence programs. Magnetic resonance in medicine. 2020;83(4):1277–1290. doi:10.1002/mrm.28020

32. Zwart NR, Pipe JG. Graphical programming interface: A development environment for MRI methods. Magnetic resonance in medicine. 2015;74(5):1449–1460. doi:10.1002/mrm.25528

33. Ravi KS, Potdar S, Poojar P, et al. Pulseq-Graphical Programming Interface: Open source visual environment for prototyping pulse sequences and integrated magnetic resonance imaging algorithm development. Magnetic resonance imaging. 2018;52:9–15. doi:10.21105/joss.01725

34. Herz K, Mueller S, Perlman O, et al. Pulseq-CEST: Towards multi-site multi-vendor compatibility and reproducibility of CEST experiments using an open-source sequence standard. Magnetic resonance in medicine. 2021;86(4):1845–1858. doi:10.1002/mrm.28825

35. Tong G, Gaspar AS, Qian E, et al. A framework for validating open-source pulse sequences. Magnetic resonance imaging. 2021;87:7–18. doi:10.1016/j.mri.2021.11.014

36. Barral JK, Gudmundson E, Stikov N, Etezadi-Amoli M, Stoica P, Nishimura DG. A robust methodology for in vivo T1 mapping. Magn Reson Med. Oct 2010;64(4):1057–67. doi:10.1002/mrm.22497

37. Santos JM, Wright GA, Pauly JM. Flexible real-time magnetic resonance imaging framework. In: Conf Proc IEEE Eng Med Biol Soc. 2004:1048–1051.

38. Lee Y, Callaghan MF, Acosta-Cabronero J, Lutti A, Nagy Z. Establishing intra-and inter-vendor reproducibility of T1 relaxation time measurements with 3T MRI. Magn Reson Med. Jan 2019;81(1):454–465. doi:10.1002/mrm.27421

39. Leutritz T, Seif M, Helms G, et al. Multiparameter mapping of relaxation (R1, R2*), proton density and magnetization transfer saturation at 3 T: A multicenter dual-vendor reproducibility and repeatability study. Human brain mapping. 2020;41(15):4232–4247. doi:10.1002/hbm.25122

40. Helms G, Dathe H, Kallenberg K, Dechent P. High-resolution maps of magnetization transfer with inherent correction for RF inhomogeneity and T1 relaxation obtained from 3D FLASH MRI. Magn Reson Med. Dec 2008;60(6):1396–407. doi:10.1002/mrm.21732

41. Haase A, Frahm J, Matthaei D, Hänicke W, Merboldt KD. FLASH imaging: rapid NMR imaging using low flip-angle pulses. 1986. J Magn Reson. Dec 2011;213(2):533–41. doi:10.1016/j.jmr.2011.09.021

42. Inati SJ, Naegele JD, Zwart NR, et al. ISMRM Raw data format: A proposed standard for MRI raw datasets. Magnetic resonance in medicine. 2017;77(1):411–421. doi:10.1002/mrm.26089

43. Gorgolewski KJ, Auer T, Calhoun VD, et al. The brain imaging data structure, a format for organizing and describing outputs of neuroimaging experiments. Scientific data. 2016;3(1):1–9. doi:10.1038/sdata.2016.44

44. Karakuzu A, Appelhoff S, Auer T, et al. qMRI-BIDS: an extension to the brain imaging data structure for quantitative magnetic resonance imaging data. medRxiv. 2021:2021.10.22.21265382. doi:10.1101/2021.10.22.21265382

45. Cohen-Adad J, Alonso-Ortiz E, Abramovic M, et al. Generic acquisition protocol for quantitative MRI of the spinal cord. Nature protocols. 2021;16(10):4611–4632. doi:10.1038/s41596-021-00588-0

46. Karakuzu A, Boudreau M, Duval T, et al. qMRLab: Quantitative MRI analysis, under one umbrella. Journal of Open Source Software. 2020;5(53):2343. doi:10.21105/joss.02343

47. Avants BB, Tustison NJ, Song G, Cook PA, Klein A, Gee JC. A reproducible evaluation of ANTs similarity metric performance in brain image registration. Neuroimage. 2011;54(3):2033–2044. doi:10.1016/j.neuroimage.2010.09.025

48. Jenkinson M, Beckmann CF, Behrens TEJ, Woolrich MW, Smith SM. FSL. Neuroimage. 2012;62(2):782–790. doi:10.1016/j.neuroimage.2011.09.015

49. Keenan KE, Gimbutas Z, Dienstfrey A, Stupic KF. Assessing effects of scanner upgrades for clinical studies. Journal of Magnetic Resonance Imaging. 2019;50(6):1948–1954. doi:10.1002/jmri.26785

50. Rousselet GA, Pernet CR, Wilcox RR. Beyond differences in means: robust graphical methods to compare two groups in neuroscience. European Journal of Neuroscience. 2017;46(2):1738–1748. doi:10.1111/ejn.13610

51. Harrell FE, Davis CE. A new distribution-free quantile estimator. Biometrika. 1982;69(3):635–640. doi:10.2307/2335999

52. Karakuzu A, Boudreau M, Cohen-Adad J, Stikov N. Thinking outside the blackbox: A fully transparent T1 mapping pipeline. In: Proc. Intl. Soc. Mag. Reson. Med. 28 (2020). 2020:4791.

53. Karakuzu A, Boudreau M, Duval T, et al. The qMRLab workflow: From acquisition to publication. In: Proc. Intl. Soc. Mag. Reson. Med. 27 (2019). 2019:4832.

54. Hansen MS, Sørensen TS. Gadgetron: an open source framework for medical image reconstruction. Magnetic resonance in medicine. 2013;69(6):1768–1776. doi:10.1002/mrm.24389

55. Assländer J, Cloos MA, Knoll F, Sodickson DK, Hennig J, Lattanzi R. Low rank alternating direction method of multipliers reconstruction for MR fingerprinting. Magnetic resonance in medicine. 2018;79(1):83–96. doi:10.1002/mrm.26639

56. Knopp T, Grosser M. MRIReco. jl: An MRI reconstruction framework written in Julia. Magnetic Resonance in Medicine. 2021;86(3):1633–1646. doi:10.1002/mrm.28792

57. Maier O, Baete SH, Fyrdahl A, et al. CG-SENSE revisited: Results from the first ISMRM reproducibility challenge. Magnetic resonance in medicine. 2021;85(4):1821–1839. doi:10.1002/mrm.28569

58. Preibisch C, Deichmann R. Influence of RF spoiling on the stability and accuracy of T1 mapping based on spoiled FLASH with varying flip angles. Magnetic Resonance in Medicine. 2009;61(1):125–135. doi:10.1002/mrm.21776

59. Heule R, Ganter C, Bieri O. Variable flip angle T1 mapping in the human brain with reduced t2 sensitivity using fast radiofrequency-spoiled gradient echo imaging. Magnetic Resonance in Medicine. 2016;75(4):1413–1422. doi:10.1002/mrm.25668

60. Ashton E. Quantitative MR in multi-center clinical trials. Journal of Magnetic Resonance Imaging. 2010;31(2):279–288. doi:10.1002/jmri.22022

61. Mancini M, Karakuzu A, Cohen-Adad J, Cercignani M, Nichols TE, Stikov N. An interactive meta-analysis of MRI biomarkers of myelin. eLife. 2020;9doi:10.7554/elife.61523

62. Lazari A, Lipp I. Can MRI measure myelin? Systematic review, qualitative assessment, and meta-analysis of studies validating microstructural imaging with myelin histology. Neuroimage. 2021:117744. doi:10.1016/j.neuroimage.2021.117744

63. Boshkovski T, Kocarev L, Cohen-Adad J, et al. The R1-weighted connectome: complementing brain networks with a myelin-sensitive measure. Network Neuroscience. 2021;5(2):358–372. doi:10.1162/netn_a_00179

64. Badji A, de la Colina AN, Boshkovski T, et al. A Cross-Sectional Study on the Impact of Arterial Stiffness on the Corpus Callosum, a Key White Matter Tract Implicated in Alzheimer’s Disease. Journal of Alzheimer’s Disease. 2020;77:591–605. doi:10.3233/JAD-200668

65. Badji A, Noriega de la Colina A, Karakuzu A, et al. Arterial stiffness cut-off value and white matter integrity in the elderly. NeuroImage: Clinical. 2020/01/01/ 2020;26:102007. doi:10.1016/j.nicl.2019.102007

66. Weiskopf N, Edwards LJ, Helms G, Mohammadi S, Kirilina E. Quantitative magnetic resonance imaging of brain anatomy and in vivo histology. Nature Reviews Physics. 2021/08/01 2021;3(8):570–588. doi:10.1038/s42254-021-00326-1

67. Salah K, Rehman MHU, Nizamuddin N, Al-Fuqaha A. Blockchain for AI: Review and Open Research Challenges. IEEE Access. 2019;7:10127–10149. doi:10.1109/ACCESS.2018.2890507

68. Moritz M, Redlich T, Günyar S, Winter L, Wulfsberg JP. On the economic value of open source hardware–case study of an open source magnetic resonance imaging scanner. Journal of Open Hardware. 2019;3(1):2. doi:10.5334/joh.14

69. DuPre E, Holdgraf C, Karakuzu A, et al. Beyond advertising: New infrastructures for publishing integrated research objects. PLOS Computational Biology. 2022;18(1):e1009651. doi:10.1371/journal.pcbi.1009651

70. Yarnykh VL. Actual flip-angle imaging in the pulsed steady state: A method for rapid three-dimensional mapping of the transmitted radiofrequency field. Magnetic Resonance in Medicine. 2007;57(1):192–200. doi:10.1002/mrm.21120

71. Gopalan K, Tamir JI, Arias AC, Lustig M. Quantitative anatomy mimicking slice phantoms. Magnetic Resonance in Medicine. 2021;86(2):1159–1166. doi:10.1002/mrm.28740

